# Reproductive biology of wild and domesticated *Ensete ventricosum*: Further evidence for maintenance of sexual reproductive capacity in a vegetatively propagated perennial crop

**DOI:** 10.1101/2020.04.30.055582

**Authors:** Solomon Tamrat, James S. Borrell, Eleni Shiferaw, Tigist Wondimu, Simon Kallow, Rachael M. Davies, John B. Dickie, Gizachew W. Nuraga, Oliver White, Feleke Woldeyes, Sebsebe Demissew, Paul Wilkin

**Author notes:** Denotes joint first authors (Solomon Tamrat, James S. Borrell).

## Abstract

- Loss of sexual reproductive capacity has been proposed as a syndrome of domestication in vegetatively propagated crops, but there are relatively few examples from agricultural systems. In this study we compare sexual reproductive capacity in wild (sexual) and domesticated (vegetative) populations of enset (*Ensete ventricosum* (Welw.) Cheesman), a tropical banana relative and Ethiopian food security crop.
- We examined floral and seed morphology and germination ecology across 35 wild and domesticated enset. We surveyed variation in floral and seed traits, including seed weight, viability and internal morphology, and germinated seeds across a range of constant and alternating temperature regimes to characterise optimum germination requirements.
- We report highly consistent floral allometry, seed viability, internal morphology and days to germination in wild and domesticated enset. However, seeds from domesticated plants responded to cooler temperatures with greater diurnal range. Shifts in germination behaviour appear concordant with a climatic envelope shift in the domesticated distribution.
- Our findings provide evidence that sexual reproductive capacity has been maintained despite long-term near-exclusive vegetative propagation in domesticated enset. Furthermore, certain traits such as germination behaviour and floral morphology, may be under continued selection, presumably through rare sexually reproductive events. Compared to sexually propagated crops banked as seeds, vegetative crop diversity is typically conserved in living collections that are more costly and insecure. Improved understanding of sexual propagation in vegetative crops may have applications in germplasm conservation and plant breeding.

## Introduction

The erosion of plant genetic resources poses a substantial threat to food security and the diverse benefits we derive from useful plants (Borrell et al., 2019; Powell et al., 2018). To address this challenge a significant effort has been made to conserve >2 million unique accessions, representing >16,500 plant species in 1750 gene banks worldwide (Commission for genetic resources on food and agriculture 2010; Fu 2017). However an emphasis on conventional seed crops may overlook the majority of perennial fruit crops for which vegetative propagation is the predominant means of replication and seed generation may be rare or absent (Miller and Gross 2011; Castañeda-álvarez *et al.* 2016; Migicovsky and Myles 2017). Vegetatively propagated crops are especially important in the tropics (Denham *et al.* 2020), where *ex situ* or *in vitro* germplasm collections are currently the only viable approach for conserving genetic resources (Thormann and Dulloo 2006). Maintaining such collections is often logistically challenging and prohibitively expensive, particularly for developing countries (Dulloo *et al.* 2013). Improving our understanding of sexual reproduction in vegetatively propagated perennial crops, for example through sexually reproducing wild progenitors (Miller and Gross 2011), has potential to address these challenges by enabling a broader range of useful plants to be banked as seed and used in breeding programmes, enhancing conservation and use of genetic diversity (McKey *et al.* 2010; J. S. Borrell *et al.* 2019; Pironon *et al.* 2019; Denham *et al.* 2020).

Clonally propagated food crops encompass at least 34 families and a wide variety of morphological diversity (McKey *et al.* 2010). This diversity has hindered attempts to define a domestication syndrome (McKey *et al.* 2010; Miller and Gross 2011; Denham *et al.* 2020), but one commonality is the hypothesis that prolonged vegetative reproduction can lead to the loss of sexual reproductive capacity (Eckert 2002; McKey *et al.* 2010; Barrett 2015; Denham *et al.* 2020). For example, domestication of Pineapple (*Annas*) which involved both sexual and asexual selection, resulted in reduced seed production through lower fertility and self-incompatibility (Chen *et al.* 2019). Loss of sexual reproductive capacity, if widespread across domesticated vegetative crops, could significantly hinder future crop breeding programmes and integration of useful alleles from crop wild relatives (Dempewolf *et al.* 2014; Migicovsky and Myles 2017). Importantly, sexual reproductive capacity does not have to be completely lost – reduced fertility, viability, altered floral allometry or germination behaviour, as a result of deleterious mutations and genetic drift in reproductive traits, could still significantly hinder programmes that seek to recombine or conserve diversity as seeds (McClure *et al.* 2014; Iriondo *et al.* 2017; Migicovsky and Myles 2017; Munguía-Rosas and Jácome-Flores 2020). However, surprisingly few studies have attempted to survey the sexual reproductive capacity of vegetatively propagated crops in agricultural systems (Elias *et al.* 2001; Scarcelli *et al.* 2006).

Here, we investigate the impact of domestication on the reproductive biology of the perennial food security crop enset (*Ensete ventricosum* (Welw) Cheesman) (Fig. 1). Enset is a giant monocarpic herb, in the sister genus to the more widely known bananas (*Musa* L.), which provides a staple starch source for 20 million people of South and South West Ethiopia (Borrell, Biswas, *et al.* 2019). Enset is a useful system in which to survey maintenance of sexual reproductive potential, because wild enset reproduction is exclusively sexual, whilst partly sympatric domesticated enset is exclusively propagated vegetatively (Borrell et al., 2020) (Fig. 1B, 1C). The latter is achieved through removal of the apical meristem and meristematic tissue of a two to three year old corm, resulting in the generation of adventitious buds (Karlsson, Dalbato, Tamado, & Mikias, 2014). Domesticated enset is not currently banked as seed in international (or national) collections (Guzzon and Müller 2016), and only a handful of local institutes maintain field germplasm collections (specifically; Areka, Yerefezy, Angacha and Hawassa, situated in southern Ethiopia). With increasing pressures due to climate change (Conway and Schipper 2011) and emerging pests and pathogens (Blomme *et al.* 2017), this represents a significant risk for the future sustainability of enset agriculture.

**Fig. 1.**
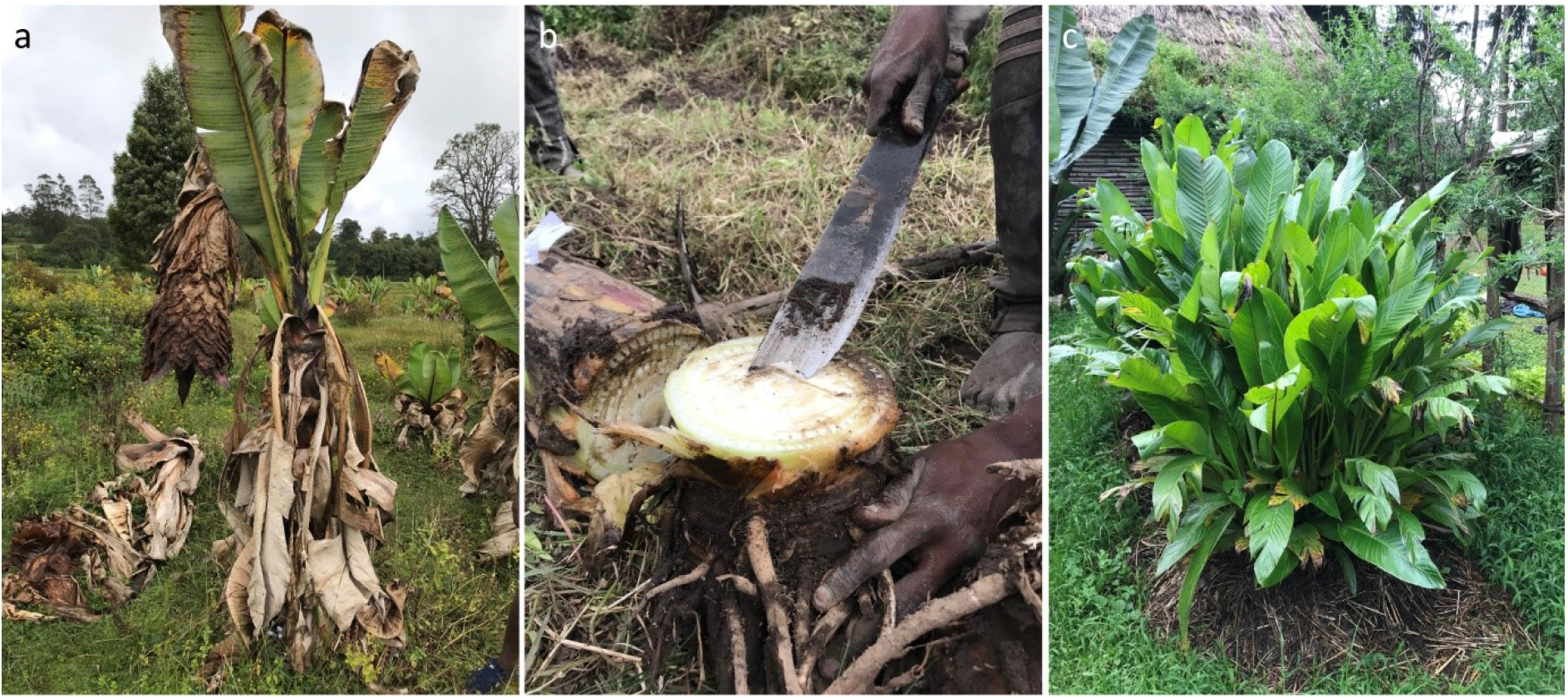
Enset cultivation in Ethiopia. A) Mature enset flowering in a neglected field near Checha. B) A farmer removing the meristematic tissue of a two-year old enset, as part of processing for vegetative propagation. C) Numerous adventitious buds sprouting from prepared corm, near Bonga.

Conserving domesticated enset diversity as seeds has been considered challenging for several reasons. Firstly, enset is monocarpic and is harvested before flowering (unlike banana), to avoid reallocation of resources from edible storage organs to the inedible inflorescence (Borrell et al., 2020). This means that developed flowers and fruits are rarely encountered in cultivation. Second, previous studies have found enset germination to be highly variable (0-90% success) (Tesfaye 1992; Messele 1994; Diro *et al.* 2003). The most detailed study to date by Karlsson et al (2013) found 5-55% germination depending on the accession. Third, recent genomic analysis concluded that accumulation of mutations in genes associated with flower initiation and seed development may have contributed to enset’s domestication (Tesfamicael *et al.* 2020). Finally, unlike other tuberous perennials, such as Yam (*Dioscorea* L.) (Mengesha *et al.* 2013) or Cassava (*Manihot* Mill.) (Rival and Mckey 2008), wild or sexually reproduced volunteer seedlings are not (knowingly) incorporated into cultivated populations (Borrell et al., 2020). Therefore, little to no indigenous knowledge pertains to enset sexual reproduction or seed germination.

In this study, we hypothesise that if sexual reproductive capacity has been conserved through domestication, we should observe consistent floral trait allometry and comparable seed morphology, viability and germination behaviour between wild and domesticated enset plants (see for example maintenance of silicon herbivore-defence through domestication in grasses, Simpson et al., 2017), or potentially, moderately divergent selection for the domesticated environmental niche (Meyer *et al.* 2012). An alternative hypothesis would be that, after being released from selection pressure, enset reproductive traits would be susceptible to deleterious somatic mutations and genetic drift. Therefore, traits would be expected to display higher variance in domesticated individuals than wild plants. A third potential scenario entails a bottleneck (though these are likely weaker in perennial vegetative crops (Gaut *et al.* 2015)) in which domesticated enset represents a subset of wild diversity, and therefore a subset of the morphological or behavioural diversity in reproductive traits. To address these scenarios, we frame our analysis with three questions; i) Is there evidence for changes in floral trait allometry, seed morphology or viability in domesticated compared to wild enset? ii) do domesticated enset seed and floral traits exhibit higher variance than under neutral expectations? iii) Does seed germination behaviour differ between wild and domesticated enset, and how do these germination requirements relate to the climate of wild and domesticated enset distributions? Finally, we discuss implications for maintenance of enset germplasm resources.

## Materials and Methods

### Sample collection

Enset in Ethiopia is readily distinguished in the field as the only member of its genus and may be subdivided into i) domesticated clonal landraces with farmers’ vernacular names, and ii) wild sexually reproducing populations. We made 20 seed collections from seven wild and 13 domesticated individuals, in Spring 2018 (Table 1). We also separately collected 15 complete inflorescences comprising six wild and nine domesticated individuals (Table 2). Domesticated accessions were collected with permission from farmers’ fields, and wild collections were made in river valleys at least 1 km from settlements cultivating enset, to mitigate the risk of feral or recently introgressed individuals. We sought to minimise collection distance between accessions to minimise differences arising from the maternal climatic environment, but due to only partly overlapping wild and domesticated distributions and the rarity of flowering individuals at the appropriate stage of maturity we highlight that our disjunct sampling may introduce additional variance.

**Table 1.** Wild and domesticated enset accessions sampled for seed morphology and germination analysis.

| Origin | Landrace | Elevation<br>(m a.s.l.) | Latitude<br>(°N) | Longitude<br>(°E) | Tetrazolium<br>viability (%) | Annual<br>Mean<br>Temp.<br>(°C) | Mean<br>Diurnal<br>Range<br>(°C) | Seed<br>count |
| --- | --- | --- | --- | --- | --- | --- | --- | --- |
| Domesticated | Ganticho | 1840 | 6.47 | 38.35 | 30 | 18.41 | 14.25 | 100 |
| Domesticated | Midasho | 2786 | 6.47 | 38.54 | 14 | 12.89 | 13.67 | 7 |
| Domesticated | Kiticho | 2787 | 6.48 | 38.54 | 50 | 12.89 | 13.67 | 16 |
| Domesticated | Gefetano | 1930 | 6.83 | 37.75 | 75 | 20.24 | 14.72 | 300 |
| Domesticated | Wanadiya | 2125 | 6.86 | 37.79 | 48 | 18.78 | 14.44 | 530 |
| Domesticated | Gefetano | 2125 | 6.86 | 37.79 | 46 | 18.78 | 14.44 | 540 |
| Domesticated | Maze | 2120 | 6.87 | 37.79 | 64 | 18.78 | 14.44 | 1265 |
| Domesticated | Hala | 2120 | 6.87 | 37.79 | 66 | 18.78 | 14.44 | 890 |
| Domesticated | Suitiya | 2120 | 6.87 | 37.79 | 0 | 18.78 | 14.44 | 75 |
| Domesticated | Deri'ea | 2700 | 7.93 | 37.9 | 0 | 15.37 | 12.97 | 600 |
| Domesticated | Lemat | 2053 | 8.45 | 38.03 | 67 | 17.97 | 13.49 | 100 |
| Domesticated | Addis Ababa | 2420 | 9.02 | 38.78 | 100 | 16.02 | 13.25 | 55 |
| Domesticated | Addis Ababa | 2430 | 9.03 | 38.76 | 84 | 16.02 | 13.25 | 1570 |
| Wild | W1 | 1880 | 7.17 | 36.22 | 0 | 18.78 | 14.35 | 450 |
| Wild | W2 | 1850 | 7.16 | 36.2 | 38 | 18.47 | 14.3 | 135 |
| Wild | W3 | 1936 | 7.16 | 36.2 | 54 | 18.47 | 14.3 | 600 |
| Wild | W4 | 1936 | 7.16 | 36.2 | 68 | 18.47 | 14.3 | 490 |
| Wild | W5 | 1927 | 7.16 | 36.2 | 78 | 18.47 | 14.3 | 700 |
| Wild | W6 | 1936 | 7.19 | 36.2 | 52 | 18.11 | 14.14 | 680 |
| Wild | W7 | 1930 | 7.29 | 36.14 | 100 | 17.97 | 13.99 | 820 |
| Wild (non-Ethiopian) | 'Rare Palm Seeds' | - | - | - | 94 | - | - | 1000 |

**Table 2.** Wild and domesticated enset accessions sampled for floral morphology analysis

| Origin | Landrace | Elevation<br>(m a.s.l.) | Latitude<br>(°N) | Longitude<br>(°E) |
| --- | --- | --- | --- | --- |
| Domesticated | Ado | 1846 | 6.46 | 38.36 |
| Domesticated | Ganticho | 1845 | 6.46 | 38.35 |
| Domesticated | Ganticho | 1856 | 6.46 | 38.35 |
| Domesticated | Ganticho | 1833 | 6.46 | 38.35 |
| Domesticated | Midasho | 1771 | 6.46 | 38.34 |
| Domesticated | Midasho | 1853 | 6.46 | 38.35 |
| Domesticated | Midasho | 1847 | 6.46 | 38.35 |
| Domesticated | Ado | 1826 | 6.47 | 38.35 |
| Domesticated | Ado | 1781 | 6.47 | 38.35 |
| Wild | NA | 1950 | 7.15 | 36.21 |
| Wild | NA | 1900 | 7.15 | 36.21 |
| Wild | NA | 1855 | 7.17 | 36.21 |
| Wild | NA | 1855 | 7.17 | 36.21 |
| Wild | NA | 2200 | 7.12 | 35.75 |
| Wild | NA | 2200 | 7.12 | 35.75 |

To ascertain whether propensity to flower differs across landraces (enset domestication is hypothesised to select for delayed maturation, Borrell *et al.* 2020), we recorded i) vernacular names of all observed and documented landraces during six field visits (2017-2020), and ii) those observed flowering. We ranked both lists by the frequency at which landraces were observed and compared ranks using a Wilcoxon rank sum test to ascertain whether landraces observed flowering are a random sample of all observed landraces. All analyses were conducted in R software (R Core Team 2017).

### Floral trait variance and allometry

Five male flowers from each of five consecutive bracts, and where available, five female flowers, were sampled from each inflorescence and preserved in 70% ethanol. We recorded the position of flowers on the peduncle (i.e. the number of bracts and number of rows of flowers from the base of the inflorescence). Samples were carefully dissected, and morphological characters recorded using digital Vernier callipers, accurate to 0.01mm. For the female flowers, we recorded the length and width of the pedicel, pistil outer and inner tepals, style and staminodes. For the male flowers, we recorded the length of the styloid, pedicel, the five anthers and filaments and the length and width of the outer and inner tepals. Where specific tissues were absent and we were certain that this was not due to a damaged sample (for example, four instead of five filaments), this trait was recorded as zero.

To investigate morphological variation, we first applied Spearman’s rank nonparametric correlation to assess pairwise relationships for floral traits within domesticated and wild enset. We report pairwise tables of correlation coefficients. Second, we tested for a significant difference between domesticated and wild trait values. We aggregated measurements by individual plant to mitigate pseudo-replication, and then performed a Bartlett test for homogeneity of variances and an unpaired t-test for each trait, applying a Holm correction for multiple tests. All analysis were performed in R software v3.4.1 (R Core Team 2017). Finally to assess differentiation in overall floral morphology between wild and domesticated enset we used redundancy analysis (‘rda’) implemented in the R package Vegan (Oksanen *et al.* 2019). We plotted data aggregated by each individual and applied permutational multivariate analysis of variation using distance matrices (PERMANOVA) using the function ‘adonis’, to test the degree to which variance in floral traits can be explained by individual or group (i.e. domesticated or wild plants).

### Seed morphology and viability

Seeds were extracted from ripe fruits by hand and the pulp containing seeds was washed thoroughly until all the flesh was removed. After extraction and cleaning, seeds were air dried at room temperature (∼20°C) for one week, packaged and transferred to the Millennium Seed Bank, UK where they were stored at a constant 60% relative humidity for 1-2 weeks (approximate humidity of collection region). In addition to field collected enset seed, we also used non-Ethiopian enset seeds legally sourced from the online horticultural distributor Rare Palm Seeds (https://www.rarepalmseeds.com/) which are available in large quantities (hereafter RPS), permitting us to screen a wider range of experimental conditions.

For each accession, 20-50 seeds were weighed individually, using a balance accurate to 0.001g. Seed size (diameter on an x, y, and z axis) was recorded for 10-20 seeds per accession using a digital caliper accurate to 0.01mm. Due to the highly non-uniform shape of enset seeds, with no consistent long or short axis, seed volume (estimated as the cube of the three measured axes) was used for subsequent analyses. We used a Faxitron Ultrafocus x-ray (Faxitron Bioptics, LLC), to measure total seed area (cross section), endosperm area and testa thickness (outer integument) in five seeds per accession (Figure S1). Seeds were positioned with the proximal end facing upwards, and results averaged by accession. We tested each dataset for homogeneity of variances and normality before applying an unpaired t-test to evaluate differences in population means between domesticated and wild seeds.

Concurrently, tetrazolium tests were used to detect living tissue and viable seeds following the standard protocol of Leist *et al.* (2003). Briefly, 7-60 seeds per accession (mean=55) were imbibed for 24 hours, chipped to expose the endosperm and placed in 1% buffered 2,3,5-triphenyl tetrazolium chloride for two days in the dark at 30°C. Subsequently, seeds were carefully dissected, and the embryo staining pattern recorded. Germination proportions were corrected for viability during subsequent analysis.

### Germination trials

We performed four germination experiments, with temperature ranges selected based on WorldClimV2 values for the region (Fick and Hijmans 2017). We exclude treatments involving sulphuric acid, sodium hydroxide, ammonium nitrate, sodium hypochlorite or hot water as Karlsson *et al.* (2013) reported that these had no significant positive effect on germination; and indeed, scarification and 70% ethanol had significant negative effects. To mimic natural conditions we draw on germination ecology in *Musa* (Laliberté 2016), where alternating temperature regimes are known to be a key requirement in *M. balbisiana* due to exposure of the seeds on the soil surface to alternating day and night temperatures (Stotzky *et al.* 1962; Kallow *et al.* 2021) and similar behaviour reported by Tesfaye (1992) in enset.

i. Exp 1: First, using RPS seed we screened a range of 15 constant and alternating temperature regimes to guide our experimental design for subsequent germination tests. Constant conditions comprised six regimes: 10°C, 15°C, 20°C, 25°C, 30°C and 40°C. Alternating conditions included 10-20°C of diurnal variation (12h:12h) to simulate larger temperature shifts, comprising an additional nine regimes: 20/10°C, 25/10°C, 25/15°C, 30/10°C, 30/15°C, 30/20°C, 35/20°C, 40/20°C, 40/25°C.
ii. Exp 2: Based on results from the initial screening, seeds from seven wild accessions and seven domesticated accessions (those with sufficient seeds available for full trials) were exposed to a refined range of 11 temperature regimes: 10°C, 15°C, 20°C, 25°C, 30°C, 20/10°C, 25/10°C, 25/15°C, 30/10°C, 30/15°C, 30/20°C.
iii. Exp 3: To evaluate the relative importance of an absolute shift in ambient temperature (e.g. warming due to disturbance) and regular diurnal temperature shifts (i.e. our alternating temperature regimes) we exposed a subset of a subset of seven domesticated accessions plus RPS, to five constant temperatures for three months (10°C, 15°C, 20°C, 25°C, 30°C), and then moved them to the ‘optimum’ alternating temperature (25/10°C) identified in earlier tests. Available wild seed was prioritised for Exps 1/2.
iv. Exp 4: To evaluate the extent and influence of dormancy, a subset of seven domesticated accessions plus RPS, was stratified at 10°C for three months, and then transferred to five other temperature regimes: 25°C, 25/10°C, 25/15°C, 30/20°C, 30/15°C. We compare these to aggregated data from Exp2 that was not exposed to a period of stratification.

In all germination tests seeds were placed on moist sand (300g sand, 42ml de-ionised water) and sealed in clear plastic boxes (120 × 180 mm). Boxes were then sealed in plastic bags to minimise moisture loss and contamination. Each box contained 60 seeds (except where specified), with a total of 235 unique condition x accession combinations evaluated in this study. Seed boxes were placed in the corresponding incubators with either constant temperature or 12 hour alternating temperature cycles. All treatments included a 12-hour photoperiod. Germination was defined by radicle emergence ≥2 mm, and tests were scored every 3-18 days, depending on activity, for 120 days. We calculated the number of days required for 50% of the final germination count in each experimental replicate. We aggregated these data by accession, excluding any replicate with zero germination.

To test for a significant difference between wild and domesticated germination behaviour (EXP2), we fitted polynomial regression models for the logit transformed proportion germinated against a) the daily temperature change and b) the mean temperature the replicate was exposed to. For each variable we fitted two models, the first with all accessions and the second with an additional variable grouping the data by type (wild v domesticated). We then used ANOVA to test whether grouping produced a significantly better model fit. To evaluate the role of stratification and dormancy (EXP3), we plotted the absolute temperature change between the first temperature and the mean of the second temperature of the treatments, against germination proportion and applied linear regression. The effect of a 10°C stratification treatment was compared to non-stratification using an unpaired t-test (EXP4).

To understand whether differences in germination behaviour were concordant with differences in local climate, bioclimatic data for Ethiopia was sourced from WorldClim v2 (Fick and Hijmans 2017) at 2.5 arc minute resolution (∼10 km). In the first instance, we extracted climate values for our study accessions, and tested for significant differences in Annual Mean Temperature and Mean Diurnal Range using unpaired t-tests. We then collated 472 enset localities from GBIF (GBIF.org, 2018), publications (Borrell et al., 2019; Pironon et al., 2019) and field observations, and subsampled these to a 10 km grid consistent with the environmental data layers, retaining 94 unique domesticated records and 19 unique wild records. We extracted climate data for these cells and aggregated them for domesticated and wild enset separately.

## Results

### Observations, collection, processing and storage

Summary information for the 15 floral and 20 seed accessions are reported in Tables 1 and 2. Overall we surveyed 375 male flowers and 45 female flowers and harvested >8000 seeds. During field surveys, we documented 1864 observations of 453 named landraces from across the enset growing region. In addition, we recorded 39 flowering individuals of 26 landraces. After ranking by frequency of observation we found no significant difference in rank order (W = 6251.5, p = 0.60).

### Floral trait allometry

Of 120 pairwise comparisons for male flowers, 100 were significantly correlated in domesticated and 72 significant in wild. Traits such as anther length and filament length were highly correlated with each other in both wild and domesticated flowers, whereas sepal length and stylode length were highly correlated with anther length in wild flowers, but only weakly correlated in domesticated flowers. Of 78 pairwise comparisons for female domesticated flowers, 34 were significant. Overall, we found relatively weak correlation for pedicel length with other traits, whereas Ovary length and Outer whorl fused tepal width were highly correlated with other traits. Pairwise correlation coefficients for male and female flower morphological traits are reported in Tables S1 and S2 respectively. Comparison of trait variance found little differences between wild and domesticated plants (Table 3). The only significant differentiation in trait means was for Sepal width, which was significantly wider in domesticated flowers (t = −7.7, df = 12.4, p < 0.001). With no wild female flowers, it was not possible to test for significant differences in female floral trait morphology, thus tissue means are reported in Table S3.

**Table 3.**
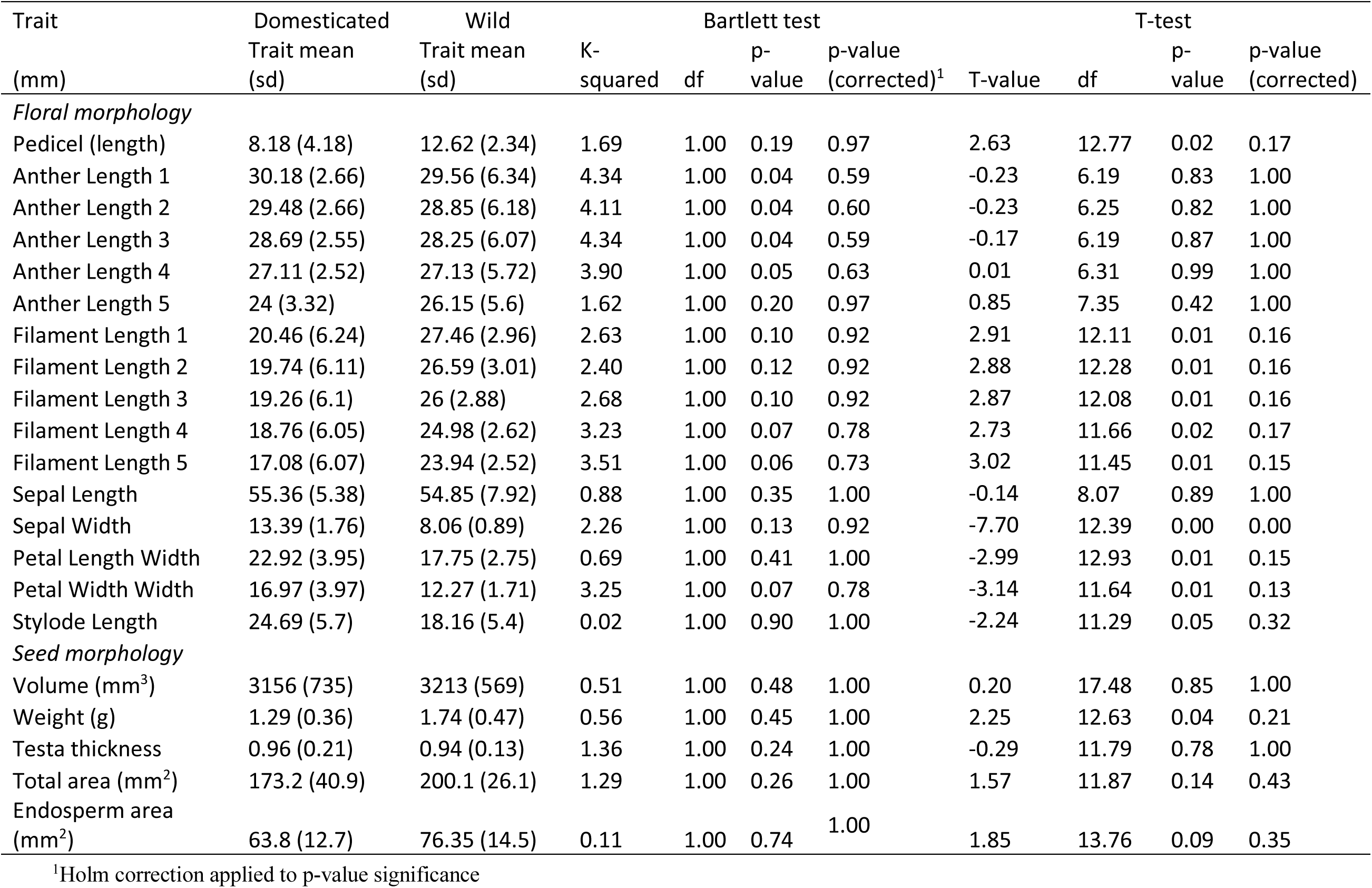
Comparison of variance and means for enset floral and seed morphology traits.

A redundancy analysis for male floral traits is plotted in Figure 2. Domesticated enset tend to vary on the first axis, with Sepal width the most important contributing variable. The majority of wild variation is on the second axis, with Filament length traits the most important contributing variables. PERMANOVA showed that the distinction between wild and domesticated origin explained a significant proportion of variation, whereas grouping samples by landrace did not (Table 4).

**Fig. 2.**
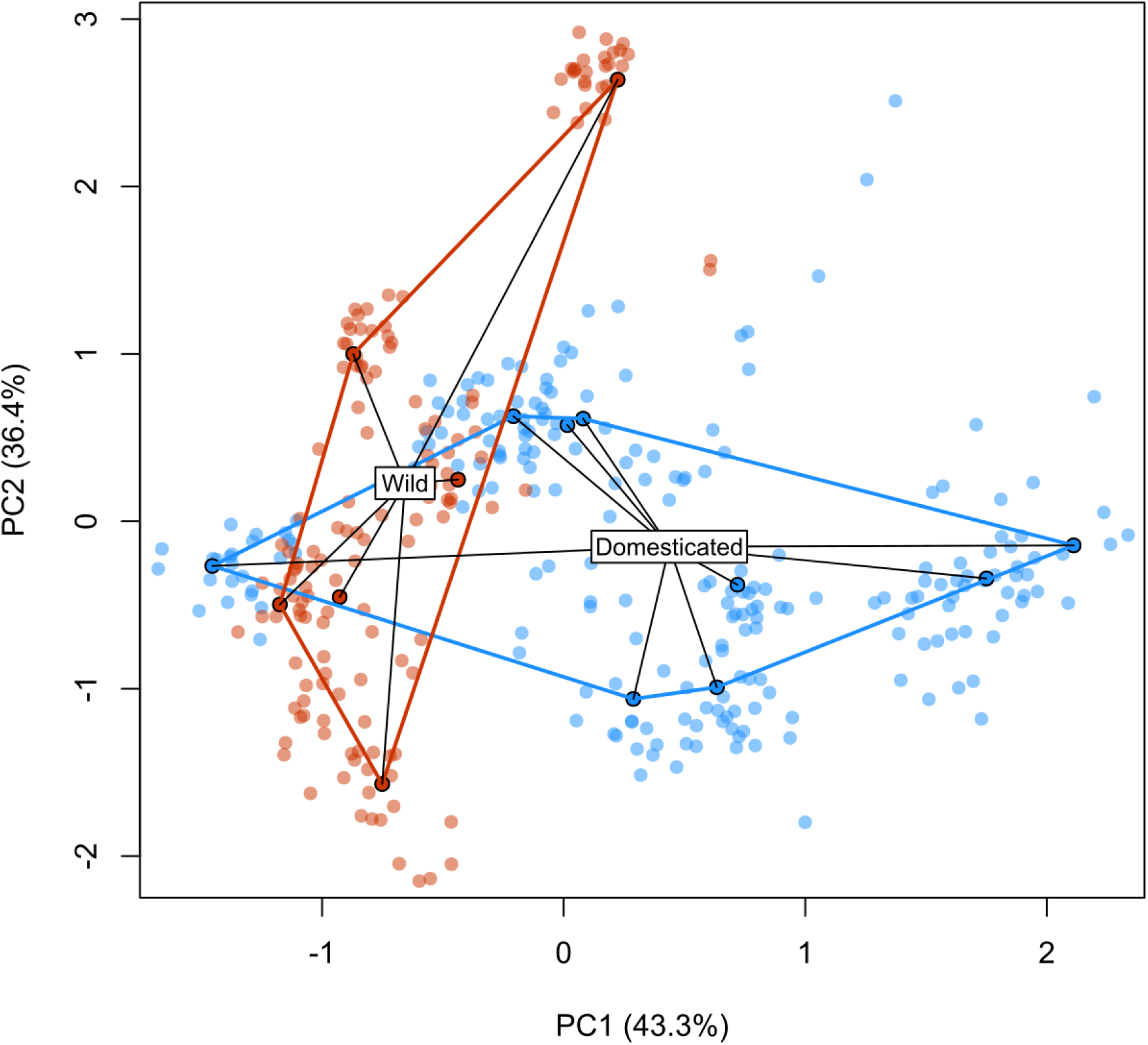
Redundancy analysis of male floral morphology in *Ensete ventricosum*. Solid points denote means aggregated by sample, background points illustrate variation of floral traits.

**Table 4.** Permutational multivariate analysis of variation using distance matrices (PERMANOVA) results for enset floral morphology with origin (domesticated vs wild) and landrace as explanatory factors.

| PERMANOVA | Df | Sums Of Sqs | Mean Sqs | F.Model | R<sup>2</sup> | Pr(>F) |
| --- | --- | --- | --- | --- | --- | --- |
| Origin (wild vs domesticated) | 1 | 49.22 | 49.22 | 3.43 | 0.23 | 0.01 |
| Landrace | 1 | 1.52 | 1.52 | 0.11 | 0.01 | 0.99 |
| Both origin and landrace | 1 | 1.36 | 1.36 | 0.09 | 0.01 | 0.99 |
| Residuals | 11 | 157.90 | 14.36 |  | 0.75 |  |
| Total | 14 | 210 |  |  | 1 |  |

### Seed morphological and viability

Seed volume ranged from 1.26 - 6.29 cm^3^ and seed weight from 0.20 g - 2.85 g in domesticated accessions, and 1.15 - 5.05 cm^3^ and 0.78 g - 2.66 g respectively in wild accessions. After correction for multiple tests we found no significant difference in seed morphology traits between wild and domesticated seeds. These patterns of (non)significance were consistent even if poorly germinating accessions are removed (*i.e.* Deri’ea, Suitiya landraces). Full morphological data are available in Tables S4 and S5. Tetrazolium tests showed high variation in viability across accessions, with both wild and domesticated accessions ranging from 0-100% viability. Mean viability was 55% and 49.5% for wild and domesticated respectively with no significant difference (t = 0.12, df = 12.9, p = 0.84) (Table 1).

### Germination trials

Mean time to 50% germination (T50) was 36 days (sd = 15.7) for domesticated enset and 35 days (sd = 8.6) for wild enset, with no significant difference detected (t = 0.51, df = 11.89, p = 0.62). In Exp 1. alternating temperature regimes outperformed constant temperatures, with the exception of constant 25°C (Figure 3). Based on these data we reduced our suite of temperature regimes in subsequent experiments. ANOVA analysis of polynomial regression models for germination behaviour in domesticated and wild enset were significantly different (i.e. grouping by accession type resulted in significantly better model fit) for both the alternating temperature range (F_136,132_ = 4.32, p = 0.003) and the mean experimental temperature (F_136,132_ = 2.71, p = 0.033) (Fig. 4A). Specifically, we found that domesticated accessions had an improved germination response in cooler mean temperatures with higher alternating temperature amplitude. Comparison of germination requirements to regional climatic conditions for wild and domesticated enset found that domesticated enset is found in local climates that have significantly cooler Annual Mean Temperatures (AMT) (t = −5.52, df = 31, p = <0.001), with significantly greater Mean Diurnal Range (MDR) (t = 3.42, df = 19.2, p = 0.003), than wild enset (Fig. 4B). Importantly, there was no significant difference in the AMT (t = −1.77, df = 12.56, p = 0.10) or MDR (t = −1.63, df = 13.96, p = 0.13) of our collected accession sites, therefore this is unlikely to be solely a maternal effect.

**Fig. 3.**
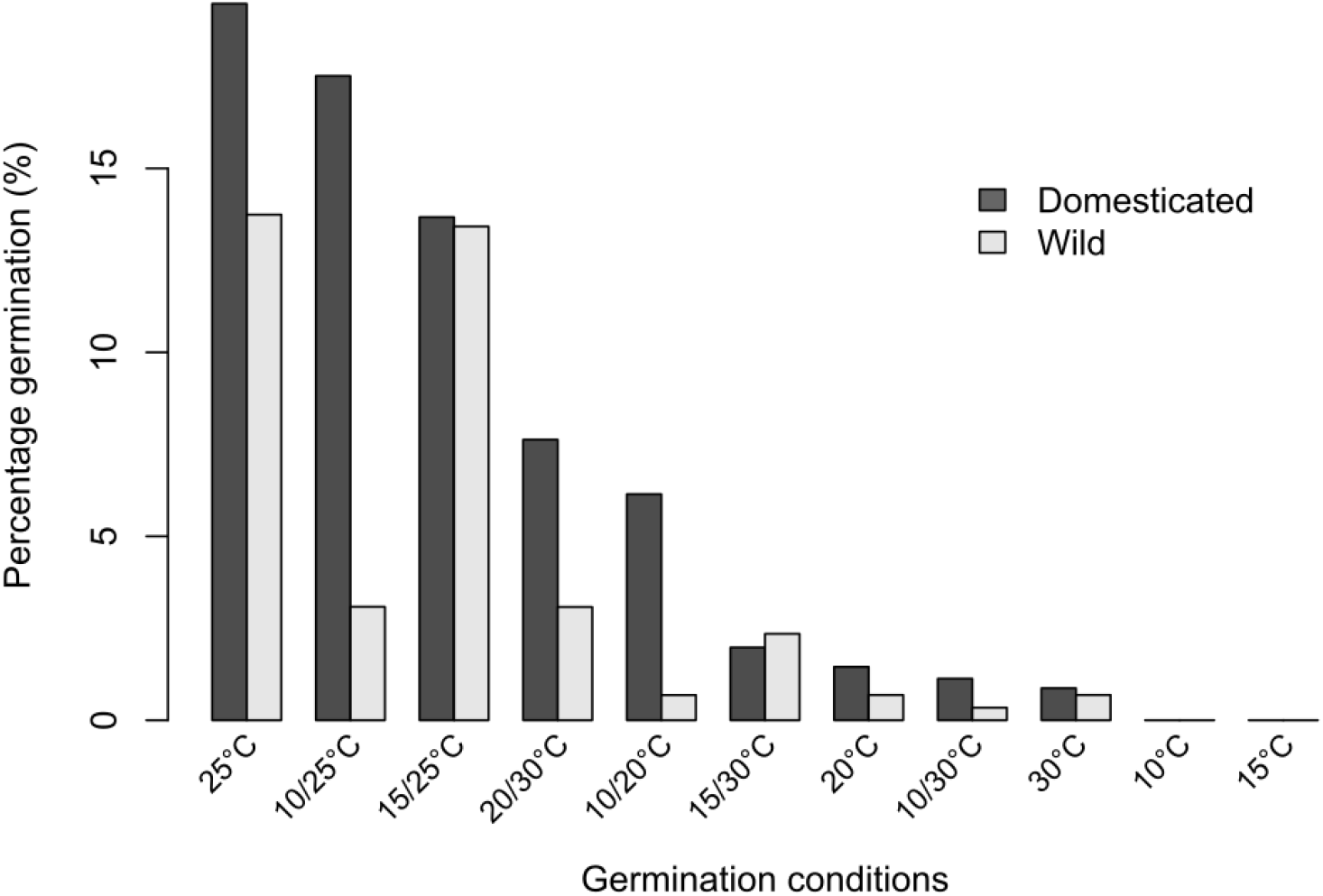
Percentage germination across a range of environmental conditions for wild and domesticated enset (Exp 2), after 120 days. Data are corrected for variation in seed viability.

**Fig. 4.**
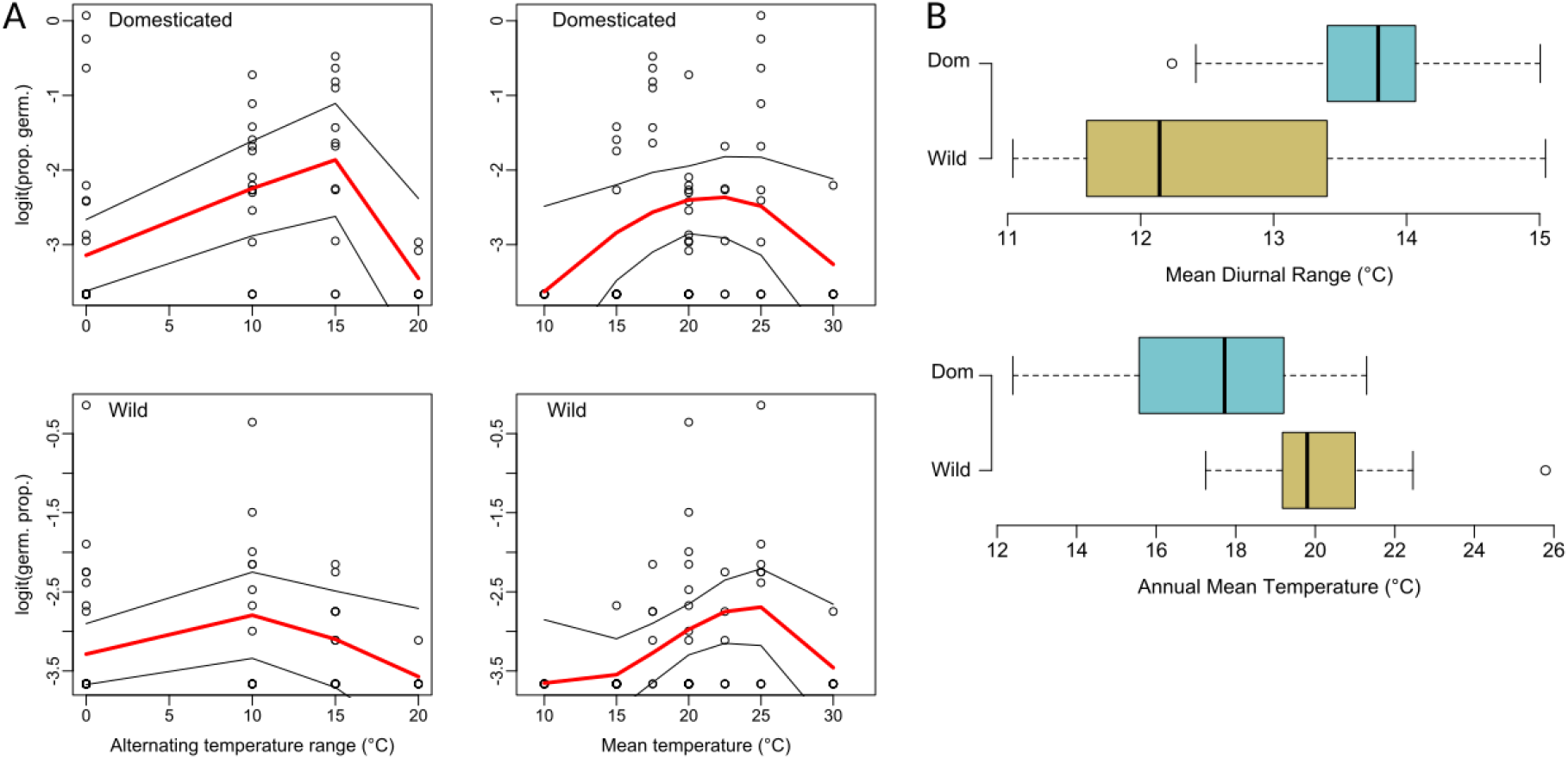
Germination behaviour and regional climate variables for wild and domesticated enset in Ethiopia. A) Polynomial regression of logit transformed germination proportion in wild and domestic accessions under varying mean temperature and alternating temperature regimes. Each point denotes a single treatment, corrected to account for variation in seed viability. B) Boxplots of regional climate for domestic and wild enset records in Ethiopia.

Analysis of Exp 3. found a significant positive relationship between germination proportion and the absolute temperature change from a constant to alternating temperature regime (F1,38 = 6.37, p = 0.0159). Analysis of Exp 4. found that a three-month period of cold stratification at 10°C prior to an experimental treatment also significantly improved germination compared to seeds immediately exposed to the experimental treatment (t = 5.22, df = 26.4, p = <0.001) (Fig. 5). Full germination data are available in Tables S6 and S7.

**Fig. 5.**
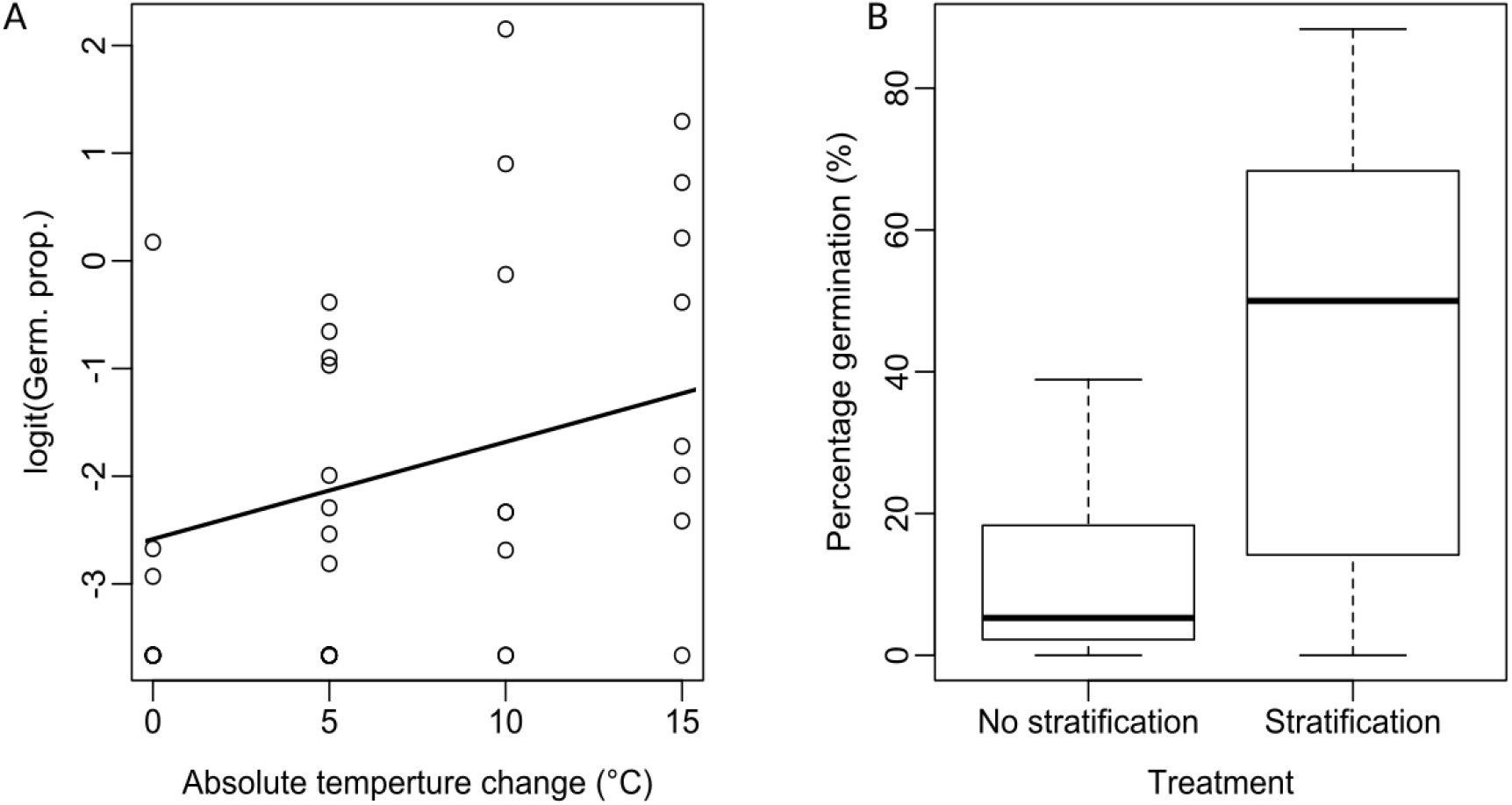
Analysis of the influence of temperature shifts and dormancy on domesticated enset germination (EXPs 3 and 4). A) The influence of absolute temperature change from constant to alternating temperature, showing a positive relationship with increased germination response for larger temperature shifts. B) Comparison of germination response for stratified seeds (3 months at 10°C), versus no stratification (immediate exposure to experimental conditions).

## Discussion

It has frequently been suggested that prolonged vegetative reproduction during domestication can lead to the loss of sexual reproductive capacity (Eckert 2002; Barrett 2015; Denham *et al.* 2020; Tesfamicael *et al.* 2020). In this study we show evidence that the indigenous Ethiopian vegetative crop enset has retained viable sexual reproductive potential through domestication. Currently, the duration over which enset has been domesticated, and the temporal advent of vegetative reproduction is unclear (Borrell *et al.* 2019). However, if we consider the extensive accumulation of indigenous knowledge associated with enset cultivation (Garedew *et al.* 2017), and its origins in the Ethiopian center of crop domestication (Harlan 1971), it is reasonable to conclude that enset was not recently domesticated. Therefore, we suggest that the potential for sexual reproduction has been maintained despite a prolonged period of vegetative propagation.

In our surveys, we found no evidence that certain landraces have lost the propensity to flower. Wild and domesticated floral morphology was significantly differentiated (figure 2), but we did not find evidence for increased variance in domesticated enset, or for loss of allometry which could be indicative of inhibited function. The exception to this pattern was sepal width which was significantly larger in domesticated flowers, even after correcting for multiple tests. It is difficult to ascertain whether this is functional, or more likely developmentally linked to another gene under selection. We found no evidence of differing seed viability rates between wild and domesticated enset, though we did find high variability in seed viability across accessions, consistent with previous reports in both enset (Karlsson *et al.* 2013) and *Musa* (Kallow *et al.* 2021). Internal and external seed morphology was also consistent between wild and domesticated accessions.

Our germination trials indicate that surprisingly, the optimum germination requirements significantly differ between domesticated and wild enset (Fig. 5). Domesticated enset has an increased germination response in cooler mean temperatures (∼22°C) and with an increased amplitude of alternating temperature (Fig. 4A) compared to wild enset. Naturally, wild enset occupies consistently warm moist tropical forest in Western Ethiopia, whereas the predominant region of contemporary enset cultivation is a region with lower Annual Mean Temperature and higher Mean Diurnal Range (Fig. 4B). This suggests, that wild and domesticated enset are potentially locally adapted to their respective environments. Importantly, there was no significant difference between the Annual Mean Temperature and Mean Diurnal Range of the wild and domesticated seed collection sites surveyed here, despite being approximately 170km apart, suggesting that this observation is unlikely to be a strong maternal effect.

When we consider enset floral morphology and seed germination behaviour together, there are several evolutionary explanations for these observations. First, a scenario where selection pressure has been relaxed as a result of domestication would be expected to show increased variance in domesticated traits, perhaps as a result of deleterious somatic mutations and genetic drift, but we do not observe this pattern in these data. If pervasive genetic drift is indeed in progress, then potentially insufficient time or generations have passed for it to become apparent. Alternatively, in a bottleneck scenario where sexual reproductive potential was not selected for, seed and floral morphology might be expected to show reduced variance, and display a subset of wild diversity. Our data do not provide strong evidence for this either.

Seeking a more parsimonious scenario, we suggest that despite virtually exclusive clonal propagation in cultivation, it is possible that a small number of escaped or neglected domesticated plants are continuing to reproduce sexually. Whilst deliberate sexual propagation is not reported by enset farmers, agricultural practices that enable sexual progeny has been recorded through ennoblement in yam (*Dioscorea*) (Cornet *et al.* 2010; Mengesha *et al.* 2013) and tolerance of volunteer manioc seedlings in Cassava (*Manihot esculenta*) (Rival and Mckey 2008). Where this has occurred in a novel agricultural environment, it is possible key functional traits have remained under balancing selection, whilst germination traits have been subjected to some degree of directional selection. This explanation is supported by both significantly different germination behaviour concordant with local environment, and lack of differences in trait means or variance. Previous work has shown that a comparatively low rate of sexual reproduction would be sufficient to maintain this balancing selection (Rice and Chippindale 2001; Cutter 2019).

More broadly, we note that enset germination requirements appear consistent with reports from *Musa* (Stotzky *et al.* 1962; Kallow *et al.* 2021). Specifically, in enset, alternating temperatures elicit a stronger germination response than constant temperatures, though in *Musa* this is virtually an absolute requirement. Using alternating temperatures as an environmental cue is hypothesised to be a strategy for detecting disturbance and canopy gaps whereby solar radiation warms the seeds in the day followed by a cooler ambient temperature at night. Enset is also reported to colonise disturbed areas (Stotzky *et al.* 1962), and thus this trait appears to be conserved across the two major branches of the Musaceae. Surprisingly, the magnitude of the transition from constant to alternating temperature was also significantly associated with germination. A possible mechanism may involve alternating temperatures reducing the ratio of Abscisic acid to Gibberellic acid (GA), reducing water potential and initiating elongation and cell growth. This suggests that future approaches involving application of GA to seeds or the growing medium may provide another mechanism for initiating germination. In addition, increased germination is also observed where seeds were stratified at a constant temperature prior to germination. Climate data indicates that the domesticated distribution of enset may reach a minimum of 8.2°C in the coolest month, which coincides with higher rainfall. Whilst no seasonality has been reported in enset flowering, this may be an additional, putatively conserved, mechanism for optimising germination timing. We anticipate that numerous factors such as fruit maturity, length and type of storage, epigenetic and other factors may influence variability, though we do not have sufficient power to resolve these in this study.

In conclusion, demonstrating the continued function of sexual reproduction in enset provides a counter example to the perceived loss of sexual capacity in vegetatively propagated crops (Eckert 2002; McKey *et al.* 2010; Barrett 2015; Denham *et al.* 2020). Development of enset seed germination protocols underpins the utility of diverse seed collections and is a compelling strategy to safeguard the genetic diversity of a food security crop, with a lower risk of provenance information loss than living plants in a germplasm collection (Thormann and Dulloo 2006). Furthermore, we provide initial evidence that very low levels of sexual reproduction may be facilitating the local adaptation of enset germination biology, influencing our interpretation of contemporary landrace diversification. An improved understanding of enset germination biology is a useful prerequisite for future crop development through sexual recombination of existing landraces, developing mapping populations and the breeding of novel genotypes, with significant reciprocal potential in bananas (*Musa*), a closely related and globally important group of clonally produced crops. In conclusion, we advocate for a broader effort in developing germination protocols for vegetatively propagated crops to provide an important alternative germplasm conservation strategy that is likely to disproportionately benefit tropical species and developing country agriculture in the global south.

## Acknowledgements

The authors thank the Ethiopian Biodiversity Institute for facilitating the export of enset seeds for germination studies. We thank John Adams and Pablo Barreiro at the Millennium Seed Bank for lab support and Dr Yann Dussert, Prof Richard Nichols, Prof Pat J.S. Heslop-Harrison and Dr Manosh Biswas for fruitful discussions on the evolutionary ecology of vegetative reproduction. We also thank Robert Kessler for permitting publication of enset photographs. We thank Mr Tobias Spanner from Rare Palm Seeds for helpful information on seed provenance and morphological variation. Finally, we thank an anonymous reviewer for helpful comments on an earlier version of this manuscript. This work was supported by the GCRF Foundation Awards for Global Agricultural and Food Systems Research, entitled, ‘Modelling and genomics resources to enhance exploitation of the sustainable and diverse Ethiopian starch crop enset and support livelihoods’ [Grant No. BB/P02307X/1]. In addition, part of this work was funded as a subgrant from the University of Queensland from the Bill & Melinda Gates Foundation project ‘BBTV mitigation: Community management in Nigeria, and screening wild banana progenitors for resistance’ [OPP1130226].

## Notes

### Competing Interest Statement

The authors have declared no competing interest.

