## Supplementary Information for "Reproductive biology of wild and domesticated *Ensete ventricosum*: Further evidence for maintenance of sexual reproductive capacity in a vegetatively propagated perennial crop"

*Denotes joint first authors

**Supplementary Tables**

**Table S1.** Male floral morphology correlation matrix

**Table S2.** Female floral morphology correlation matrix

**Table S3.** Summary of female floral trait means

**Table S4**. Seed morphology raw data – size.

**Table S5.** Seed morphology raw data – weight.

**Table S6.** Germination raw data experiments 1&2

**Table S7.** Germination raw data experiments 3&4

**Supplementary Figures**

**Figure S1.** X-ray images of *Ensete ventricosum* seeds. Scale bar denotes 5mm. A) Domesticated accession ‘Wanadiya’. A poorly filled seed can be observed in the lower center of the image. B) Wild accession six ‘W6’. C) Major enset seed morphological traits referred to in this study: 1. Hilum. 2. Micropylar plug. 3. Embryo. 4. Endosperm surrounding embryo. 5. Inner integument. 6. Outer integument (testa) 7. Chalazal mass. Scale bar denotes 5 mm.
